# Stimulus reliability automatically biases temporal integration of discrete perceptual targets

**DOI:** 10.1101/841353

**Authors:** Dragan Rangelov, Rebecca West, Jason B. Mattingley

**Affiliations:** Queensland Brain Institute, The University of Queensland, St Lucia, QLD Australia; School of Psychology, The University of Queensland, St Lucia, QLD Australia; Canadian Institute for Advanced Research (CIFAR), Toronto, Canada

## Abstract

Many decisions, from crossing a busy street to choosing a profession, require integration of discrete sensory events. Previous studies have shown that integrative decision-making favours more reliable stimuli, mimicking statistically optimal integration. It remains unclear, however, whether reliability biases are automatic or strategic. To address this issue, we asked observers to reproduce the average motion direction of two suprathreshold coherent motion signals, presented successively and varying in reliability. Although unbiased responses were both optimal and possible by virtue of task rules and suprathreshold motion coherence, we found robust behavioural biases favouring the more reliable stimulus. Using population-tuning modelling of brain activity recorded using electroencephalography, we characterised tuning to the average motion direction. In keeping with the behavioural biases, the tuning profiles also exhibited reliability biases. Taken together, our findings reveal that temporal integration of discrete sensory events is automatically and sub-optimally weighted according to stimulus reliability.

## Introduction

Decision-making is a ubiquitous cognitive process involved in many forms of choice behaviour. One pertinent issue is how multiple sources of evidence might be combined in support of a single decision. For example, to safely cross a busy street, one should consider the traffic coming from both sides of the road. While decision-making in relation to single stimuli – in our example, monitoring traffic on just one side of the street – has been well characterised at both computational [1–9] and neural [10–15] levels, much less is known about how observers integrate two (or more) distinct sources of evidence into one decision. In recent years there has been increased interest in the cognitive and neural mechanisms underpinning such complex or ‘integrative’ decision making [15–19]. Studies of integrative decisions [17,18,20–22] have shown that, when presented with two stimuli, one of higher reliability than the other, the choice behaviour of humans [22] and rodents [17] is biased in favour of the more reliable stimulus. In fact, biases in integrative decision-making closely resemble statistically optimal signal integration [23,24] in which the contributions of individual signals are weighted by their reliability so as to yield statistically optimal decisions.

While previous research has demonstrated that integrative decision-making is subject to biases based on factors such as stimulus reliability, it remains unclear whether these biases are automatic or strategic. One ubiquitous finding in the literature is that some observers do not integrate discrete sensory events at all, but rather rely exclusively on signals of higher reliability [17,18,20,22]. This finding suggests that integrated decisions may be subject to higher order influences, such as a trade-off analysis of increased accuracy at the expense of increased effort. Of particular note, studies on integrative decision-making have typically employed tasks in which the integration was helpful, but not necessary for accurate responses. While characterising higher order biases is an important open issue [25], it remains unclear whether integrated decision-making automatically favours sources of higher reliability in tasks where the integration is essential, rather than opportunistic.

To address this important issue, we developed a task that required explicit integration of two simple visual decisions. On every trial, two brief periods of coherent motion were presented and participants had to report the *average motion direction* of the two epochs, without speed constraints, by adjusting the orientation of a response dial (Fig. 1a). To illustrate, if a trial contained successive motion directions toward 10 o’clock and 2 o’clock, participants should indicate an *average* motion direction of 12 o’clock. To minimize the likelihood that participants might compute the average motion direction by simply accumulating all motions signals, on every trial we presented two overlaid patches of dots – a target patch and a distractor patch – in different colours. A colour cue at the beginning of the trial indicated the dot-patch that would contain the target-motion direction. Participants had to monitor for coherent motion events in the target patch while ignoring concurrently presented motion in the distractor patch. Throughout the task we recorded neural activity using electroencephalography to characterise direction tuning to the presented motion stimuli, as well as to observers’ internally generated integrated decisions.

**Figure 1.**
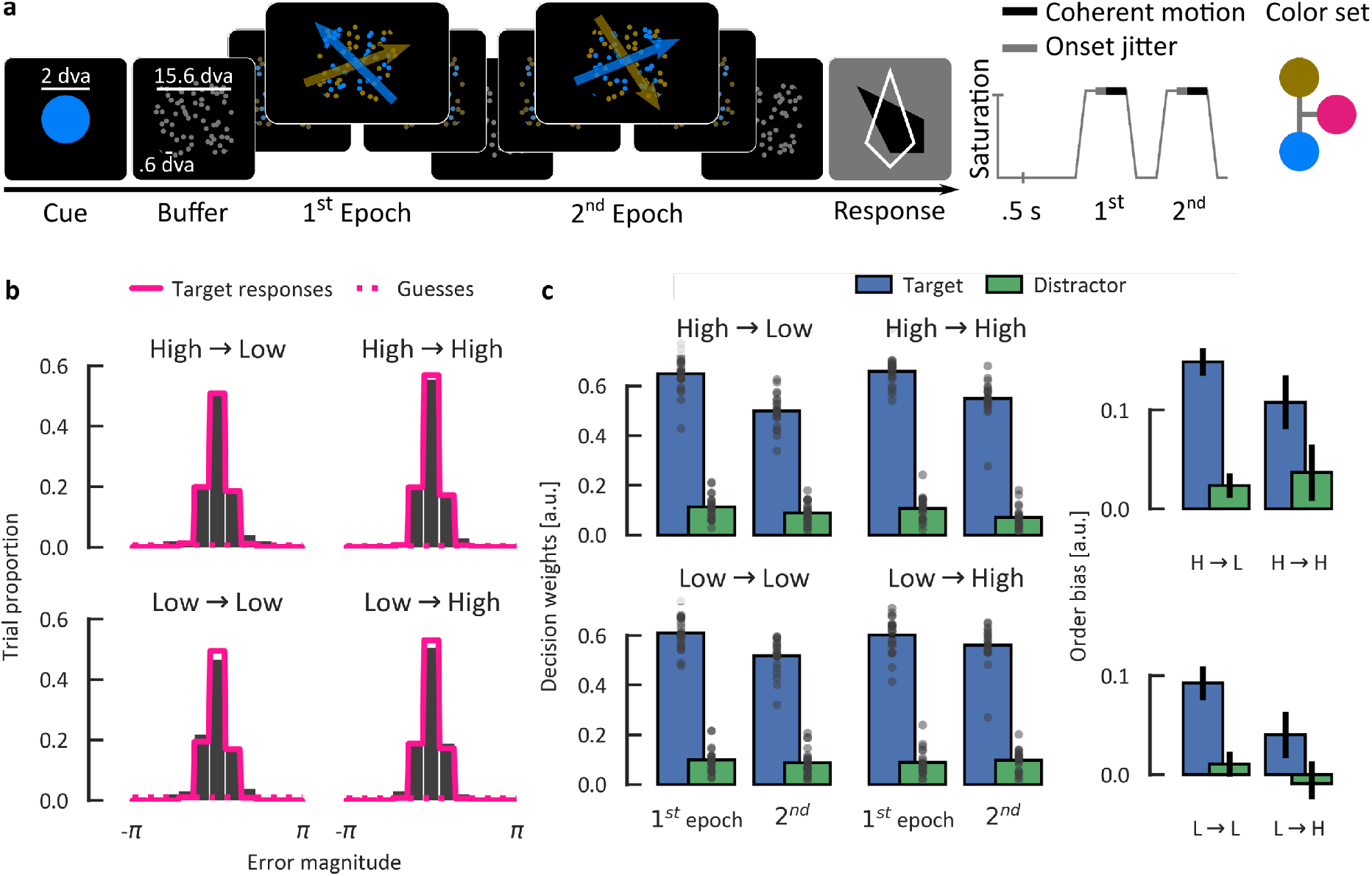
Overview of the experimental paradigm and the behavioural results. (a) A typical display sequence per trial (right) together with the time-course of different events and possible colours (left). A coloured cue indicating the task-relevant colour (fixed per participant) was followed by a patch of grey dots moving randomly. After 1 s buffer periods, the colour saturation increased gradually to reveal two intermingled fields of distinctly coloured target and distractor dots. Coherent motion signals were presented for .5 s in both fields, jittered relative to the maximum saturation onset (.25–.5 s). Two epochs were presented and participants reproduced the average motion direction of the two target motion stimuli by adjusting the orientation of the response dial (black) without speed constraints (max. 6 s). Response feedback was shown for .5 s following the response (white dial contour). (b) Distributions of the observed error magnitudes (expected/correct – actual response, bars) together with the predicted distributions (pink lines) on the basis of mixture distribution modelling [27,28]. Separate models were fitted across different combinations of motion coherence (Low/High) in the first and second epoch (First epoch coherence → Second). (b) Left: Decision weights across different experimental conditions showing the degree to which coherent motion signals presented during a trial influenced the response. Right: Difference in respective decision weights for the first and second epochs separately per condition and stimulus type. Dots represent individual participants. Error bars denote within-participants standard errors of the mean [31].

To manipulate stimulus reliability, the motion coherence in the first and second epoch could either be low (40% of coherently moving dots) or high (80%). Different combinations of stimulus reliability across two epochs were presented equally often and in a random order. To facilitate unbiased integration, we used reliability levels well above threshold levels, which are typically around 6–7% for most observers [26]. In addition, in a separate control experiment we verified that observers showed equivalent accuracy in judging the direction of *single* low-or high-coherence motion targets, thereby confirming that unbiased integration in the averaging task was, in principle, possible. It follows, therefore, that if observers are able to make optimal (i.e., unbiased) integrated decisions, the target stimuli should contribute equally to average direction judgements, irrespective of their reliability. By contrast, we found that signal reliability affected both behavioural and neural markers suggesting that integrative decision-making automatically favours sources of higher reliability even when the reliability-bias yields suboptimal decisions.

## Results

In the main experiment, we presented two successive epochs of coherent motion signals on every trial (Fig. 1a) and asked participants (N=22) to reproduce the average motion direction of two target signals, while ignoring two, concurrently presented distractor signals. Inspection of distributions of the observed error magnitudes (expected/correct response – actual response) revealed a continuous, relatively narrow, unimodal distribution in all experimental conditions (Fig. 1b, grey bars). We used mixture distribution modelling [27,28] of error magnitude to independently quantify the proportion of trials on which participants guessed randomly (Pg) and the response precision (K) on the remaining, target response trials. Model fitting separately per participant and experimental condition yielded close fits to the observed errors (pink lines). Participants guessed on approximately 10% of trials (guess rate Pg_M/SEM_ = .09/.01) and the response precision of target responses was relatively high (K = 8.16/.34, FWHM = 48°) indicating that participants were able to perform the task well.

We next analysed the effect of dot coherence in different epochs on response precision and guessing rates. The response precision was comparable for low and high coherence in the first epoch (K = 8.12 and 8.19, respectively, F < 1). By contrast, response precision was significantly worse for low than high coherence targets in the second epoch (7.61 and 8.70, respectively, F_1,21_ = 14.27, p = .001, 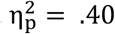). The interaction between coherence levels in the first and second epochs was not significant (F < 1). Independently of epoch, participants were more likely to guess for low coherence signals than high (.10 and .06, overall, main effect of coherence in the first epoch F_1,21_ = 25.75, p < .001, 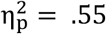, in the second F_1,21_ = 13.80, p = .001, 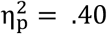, interaction F < 1).

We next ran linear regression analyses separately per participant and experimental condition using complex-valued response angles as a dependent variable and the presented motion directions as independent variables (target/distractor signals in first/second epoch; see Methods). The absolute values of the regression weights reflect how much the respective motion signals influenced integrative decision-making, i.e., their decision weights. Target weights were much larger than distractor weights (.56 and .09, respectively, F_1,21_ = 2,028, p < .001, 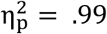, Fig. 2c), demonstrating that participants did not passively accumulate evidence from all motion signals presented in a trial. Interestingly, both target and distractor weights were higher in the first epoch than in the second (.63 and .53 for targets, respectively, F_1,21_ = 24.28, p < .001, 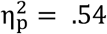; .10 and .09 for distractors, F_1,21_ = 4.60, p = .044, 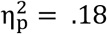). This primacy bias is opposite to what one might expect on the basis of simple memory decay [29,30], because any such decay should have affected the first (older) target more than the second (more recent) target. Rather, the primacy bias in combination with statistically significant coherence effects for the second epoch might indicate how participants represented individual signals prior to integrating them.

**Figure 2.**
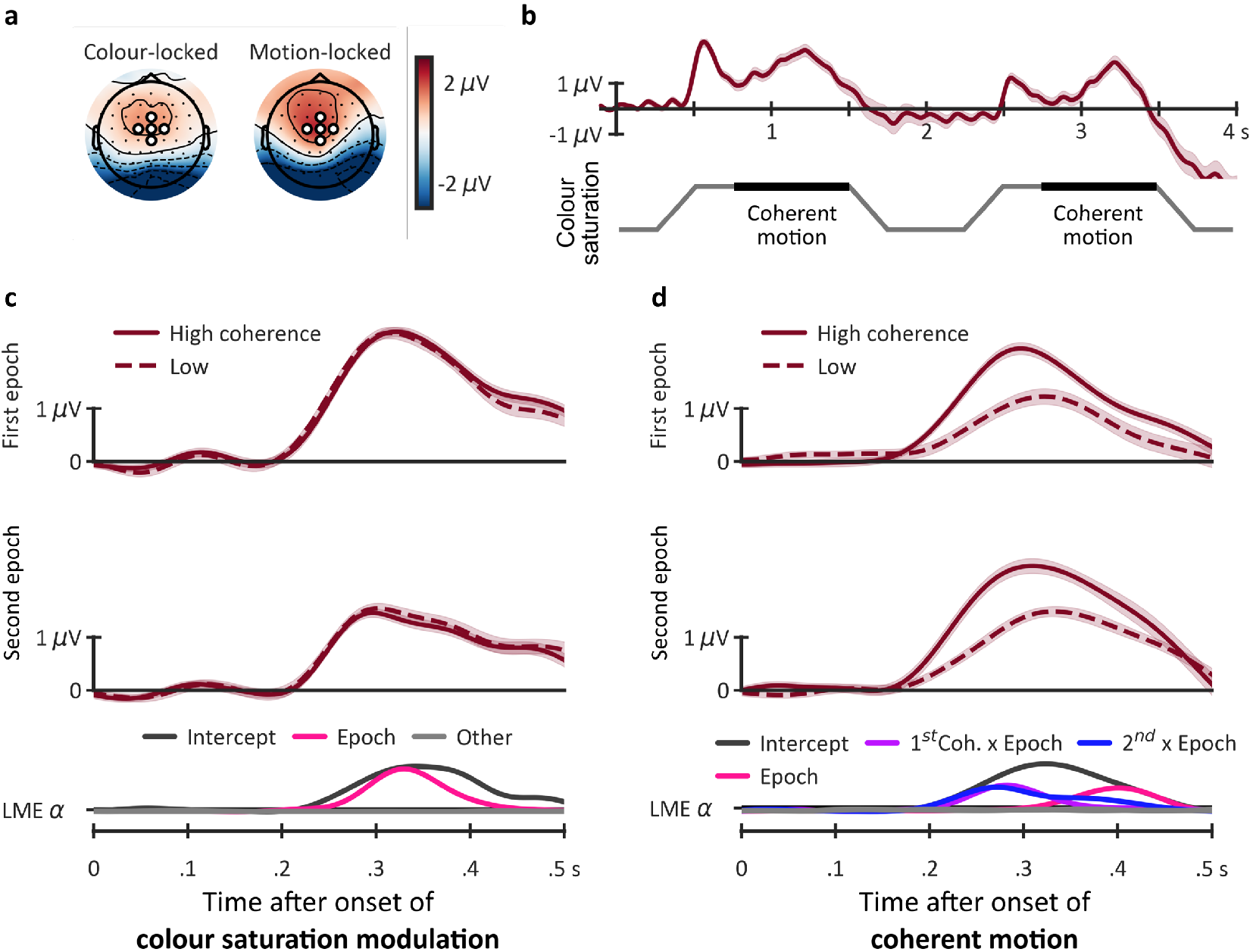
ERP results. (a) ERP topography for colour-locked responses, showing the average of time-samples at which the colour saturation reached a maximum (.75 s and 2.75 s for the first and the second epoch, respectively); and for motion-locked responses, showing the average of time-samples at which the motion signal was most likely to be presented (at 1.25 and 3.25 s). Marked electrodes (white circles) were used for computing the CPP time-course. (b) The CPP time-course (upper panel) together with the time-course of different events within a trial (lower panel). (c) Colour-locked CPP during the first .5 s following the onset of colour modulation in the first epoch (upper panel) and the second epoch (central panel). Only random motion was presented during this period. The lower panel shows −log_10_(p_FDR-corrected_) for different terms in the stepwise linear mixed effect model of colour-locked CPPs. Values above the LME a are significant at p < .05 (see Methods). (d) Motion-locked CPP. The colours were at maximum saturation during the analysed 500 ms. Conventions as in panel (b). For the purpose of presentation, all time-traces were low-pass filtered at 10 Hz using a Butterworth 4^th^ order infinite impulse-response filter. The analyses were performed on unfiltered data. Shaded areas denote ±1 within-participants SEM.

Most importantly, this primacy bias was further qualified by two-way interactions with motion coherence in both the first and second epochs (Fig 2c, right panel). Specifically, whereas increasing the coherence in the first epoch *increased* the primacy bias (F_1,21_ = 16.30, p < .001, 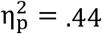), increasing the coherence in the second epoch *decreased* the bias (F_1,21_ = 4.34, p = .049, 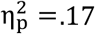). The interaction of dot coherence with target order indicates that averaging two signals is biased in favour of the signal of higher reliability. Importantly, this bias arose despite the fact that, consistent with the task instructions to ***average*** the two target signals, unbiased averaging should be the optimal response strategy. The reliability bias in decision weights suggests that integrative decision-making is automatically reliability-weighted, even when this bias leads to suboptimal decisions.

To characterise the neural correlates of integrated decision-making, we recorded brain activity using electroencephalography (EEG). We quantified a well-documented neural correlate of simple perceptual decision-making [11,12,32], the centro-parietal positivity (CPP, see Methods and Fig. 2). Visual inspection of the ERP topographies (Fig. 2a) revealed a positive deflection over centro-parietal electrodes, consistent with the typical CPP topography [33]. Visual inspection of the CPP time-course over the trial (Fig. 2b) revealed a phasic modulation that closely followed the trial sequence. During periods of grey, randomly moving dots, the CPP amplitude was at baseline. Two sharp deflections closely followed the onset of colour modulation in the first and the second epochs, and two broader deflections coincided with periods of coherent motion. A single, continuous episode of evidence accumulation should have resulted in a sustained response across the trial. Instead, the observed phasic modulation of the CPP suggests that several, temporally separable episodes of evidence accumulation took place within each trial.

We next analysed shorter segments (500 ms) of EEG data time-locked to the onsets of colour modulation and coherent motion (Fig. 2c and 2d, respectively). For both colour-and motion-locked epochs, the CPP deflection started at around 200 ms after onset, consistent with the notion that the CPP is not merely a sensory-evoked response, but rather reflects higher level processes following sensory encoding [32,33] (see the Intercept line in Fig. 2c and 2d, lower panel). The colour-locked CPP was modulated only by the epoch, with a steeper rise and higher peak amplitude in the first epoch than the second (Fig. 2c). Analyses of the motion-locked CPP revealed a robust effect of motion coherence, with a steeper rise and higher peak amplitude for high relative to low coherence in both epochs (Fig. 2d). Importantly, only the coherence value for the currently presented motion stimulus affected the CPP, as evidenced by a significant interaction between epoch (first/second) and the motion coherence per epoch (First coherence x Epoch and Second coherence x Epoch lines, lower panel). This finding suggests that the motion-locked CPP reflects evidence accumulation in support of discriminating the currently presented motion target (i.e., a simple perceptual decision), rather than the averaging process (i.e., an integrated decision).

Whereas analyses of behavioural decision weights revealed robust interactions between the primacy bias and motion coherence across the two epochs, the ERP analyses suggest that the order effect and the coherence effects might have separable neural correlates. The primacy bias, evident in the colour-locked CPP, appears to reflect the process of selecting the target patch against the distractor patch – and, potentially, staying focused on the target patch – whereas the coherence effect, evident in the motion-locked CPP, appears to reflect the strength of the subsequently presented motion signals.

We next characterised time-resolved, motion-specific neural responses to target and distractor signals (Fig. 3a) using population-tuning modelling [34–36] of the motion-locked EEG signals (see Methods). Inspection of tuning to target signals revealed a robust and sustained motion-specific response. The onset of significant motion-tuning coincided with the peak latency of the motion-locked CPP, suggesting that motion-tuning reflects the decision about the currently presented motion stimulus. There was no such tuning to distractors, even though these signals were physically identical to the targets in terms of their brightness and coherence, which further supports the notion that participants engaged in integrative decision-making, rather than passive accumulation of all presented motion signals. Importantly, tuning to the target motion direction was sustained well after motion offset (indicated by dotted vertical lines in Fig. 3), suggesting that representations of individual signals were maintained until both targets had been presented.

**Figure 3.**
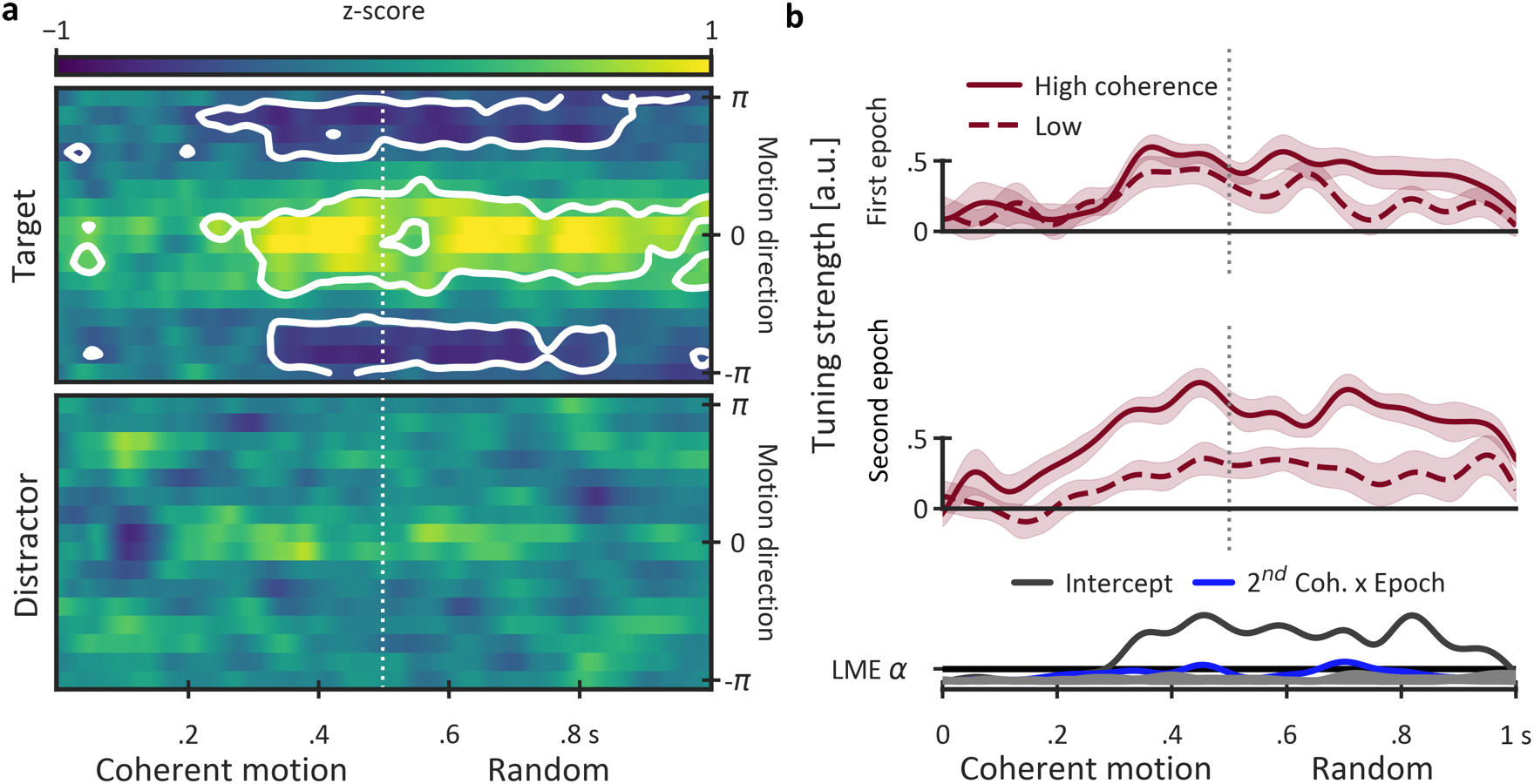
Population-tuning modelling of neural activity in response to target and distractor signals. (a) Time-resolved responses to target and distractor signals (upper and lower panels, respectively) of 16 hypothetical motion-specific channels spanning the full circle (from −π to +π) and centred on the actual presented motion signal. The trial-averaged profile of channel responses per participant and time-sample was z-scored across channels. White contours denote areas which were significantly different from 0 (FDR-corrected across all time-samples and channels). Significant modulation of channel-response profiles with yellow areas around 0 and blue areas around −π and +π indicate robust motion tuning to the presented signal. (b) Time-resolved tuning strength to *target signals* across first and second epochs (upper and lower panels, respectively). The tuning strength is an aggregate index of the channel-response profile (see Methods for details) with 0 indicating no tuning. Conventions as in Figure 2.

Inspection of the time-resolved tuning strength for *target signals* (Fig. 3b) across different epochs revealed comparable tuning to high and low coherence signals in the first epoch. This is consistent with the observation of no effect of dot coherence in the first epoch on behavioural response precision, and it supports the notion that the two coherence levels afforded decisions of comparable precision. In keeping with the significant effect of dot coherence in the second epoch on behavioural response precision, the tuning strength in the second epoch was significantly larger for high coherence than for low coherence targets. These results, in conjunction with the smaller decision weights and weaker colour-locked CPP responses in the second epoch relative to the first, suggest that the overall level of focused attention decreased from the first to the second epoch within a trial.

In a final analysis, we characterised the temporal dynamics of integrated decision-making by estimating neural tuning to the *average* motion direction (Fig. 4a). Note that this quantity is internally computed by participants, whose task was to integrate the directions of the two target signals. As such, any neural tuning to the average-target motion within a trial must reflect the individual’s integrated perceptual decision for that trial. For the first epoch, there was no significant motion tuning. This finding was expected, as neural representations of average motion direction can only be determined *after* presentation of the second motion target within the trial. By contrast, there was robust and sustained tuning to the average motion direction in the second epoch, starting from the offset of the second coherent-motion signal. Note that during this period only random motion was present on the screen, so tuning to the average motion direction could not have been stimulus-driven.

**Figure 4.**
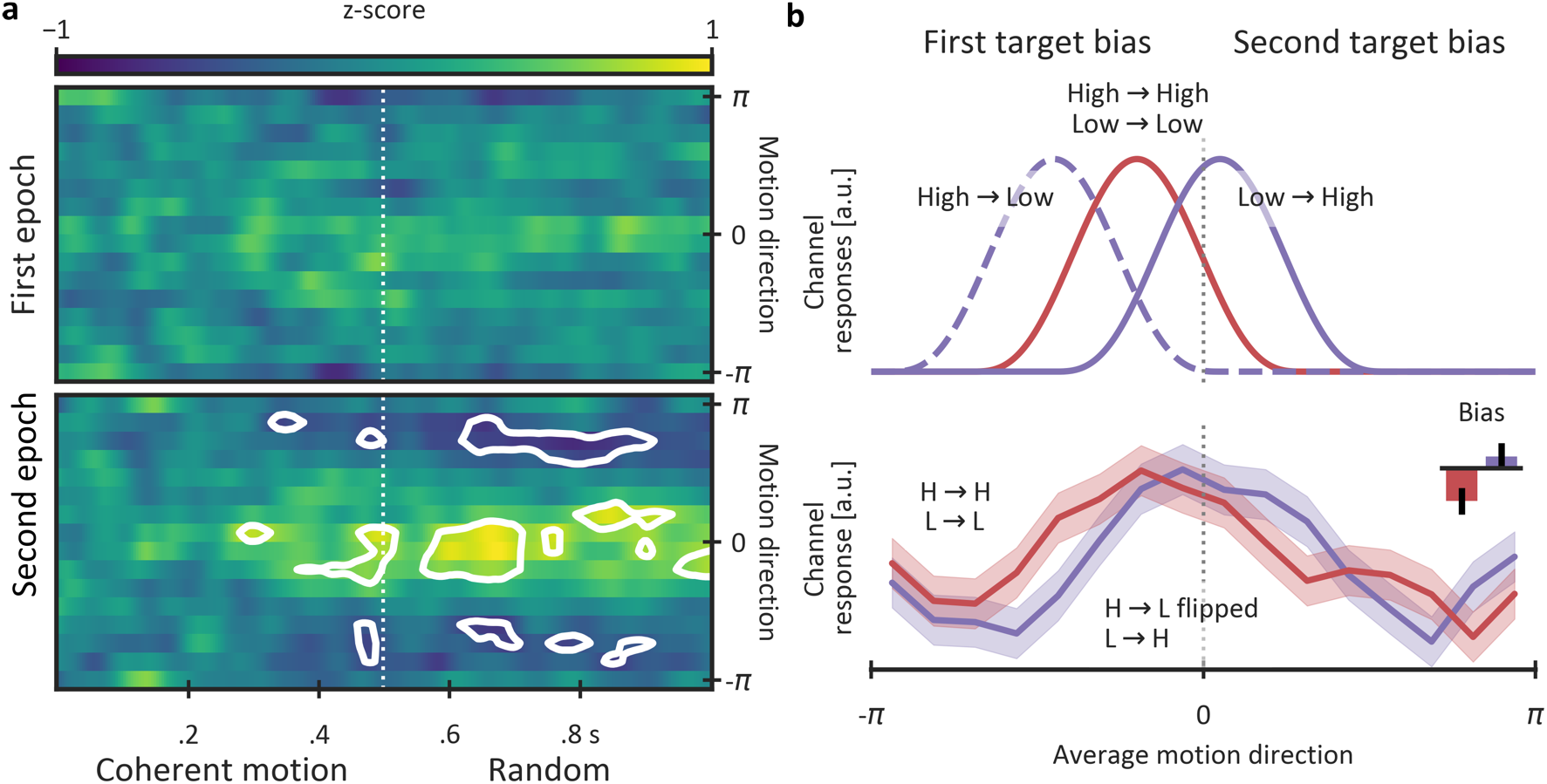
Population-tuning modelling of neural activity associated with the average motion. (a) Time-resolved average-motion tuning in the first and the second epoch (upper and lower panels, respectively). Conventions as in Figure 3. (b) *Upper panel*: Expected shifts in the tuning profile reflecting the observed behavioural order bias. The motion channels were sorted relative to the first-and the second-presented motion direction so that shifts to the left from 0 reflect a first target bias. *Lower panel*: Observed shifts in the tuning profiles for the same-coherence trials (High→High and Low→Low, red line) and different-coherence trials (High→Low and Low→High, purple line). The profiles were computed by averaging channel responses in the .5–1 s interval during which there was robust tuning to the average motion direction. To match the number of trials in the same-coherence and different-coherence trials and to maximise the signal-to-noise ratio in the different-coherence conditions, the channels were flipped for the High→Low trials so that the direction of a potential order bias would be the same as for the Low→High trials. *Small inset panel*: the average shift of the tuning profiles computed as the difference between the mean response for channels tuned to -π –0 and 0 –π intervals. Whiskers denote ±1 within-participants SEM.

To investigate how the brain integrates two discrete decisions, we next quantified the tuning profiles for the average motion direction (Fig. 4b). If stimulus reliability affects integration processes, the tuning profile should depend on the combination of motion coherence levels across target epochs within a trial. We focused on quantifying potential shifts in the profile that might reflect the first and the second target bias. To do so, the motion channels on each trial were re-coded so that negative channels were closer to the first presented target, and positive channels were closer to the second. Thus, a leftward shift would indicate a first-target (primacy) bias, and a rightward shift would indicate a second-target (recency) bias. The channels’ responses were then averaged across trials separately for different combinations of stimulus reliability across epochs.

Motivated by the first-target bias observed in behavioural decision weights (Fig. 1b), we expected to see a leftward shift for trials with the same coherence across epochs (Fig. 4b, upper panel). To increase signal-to-noise ratio, we averaged the tuning profiles for High→High and Low→Low coherence trials. The observed tuning profiles (Fig. 4b, lower panel) confirmed our expectation, as we observed a statistically significant leftward shift consistent with a first-target bias (M/SEM = -.16 /.06, t_21_ = 2.32, p_FDR-corrected_ = .046, Fig. 5b, inset panel). This result mimics the first-target bias in behavioural decision weights, and it suggests that the brain integrates two target signals in a biased way, with the first signal contributing more strongly than the second.

In contrast to the same-coherence trials (High→High, Low→Low), for the different-coherence trials we expected to observe effects of signal reliability on the tuning profile shifts. For the High→Low sequence, in which the primacy bias and the reliability bias both favoured the first target, we expected to see an even stronger leftward shift relative to the same-coherence trials (Fig. 4b, upper panel). For the Low→High sequence, on the other hand, in which the order bias and the reliability bias favoured different targets, we expected to see weaker shifts in the tuning profile relative to the same-coherence trials (Fig. 4b, upper panel). To match the numbers of trials in the same-and different-coherence conditions and to increase signal-to-noise ratio in the different-coherence condition, we flipped the tuning profiles for High→Low trials so that the expected shift direction was the same for High→Low and Low→High trials. Confirming our predictions, the first-target bias for different-coherence trials was not significantly different from zero (.05/.06, t_21_ = .66, p_FDR-corrected_ = .519, Fig. 5b, inset panel), and it was significantly smaller than the bias for same-coherence trials (t_21_ = 2.35, p_FDR-corrected_ = .046). The shifts in tuning profiles suggest that the brain relies more on signals with high reliability than low reliability when integrating two temporally separated signals.

Our findings demonstrate that integrated decisions are biased in favour of a more reliable stimulus, even though *unbiased* integration should have yielded more accurate responses in our task. The absence of a statistically significant effect of stimulus reliability in the first epoch on both behavioural response precision (K) and the tuning strength of feature-specific brain responses suggest that unbiased integration was, in principle, possible. To empirically test that the reliability levels we used in our task afforded responses of comparable precision, we conducted a control experiment in which a single epoch of coherent motion was presented and participants had to reproduce the target motion direction while ignoring a concurrently presented distractor (see Methods). Unlike in the main experiment, here participants (N=23) were shown only a single target/distractor epoch, and were thus not required to perform any motion averaging. The dot coherence varied randomly across trials between low (40%) and high (80%). We recorded behavioural responses, which we analysed using mixture distribution modelling and linear regression.

The estimated response precision was high and comparable between low and high coherence (K_M/SEM_ = 14.86/.71 and 13.73/.71, respectively, one-sample t_22_ = 1.12, p_one-tailed_ = .137). Similarly, the estimated guessing rates were very low and comparable between coherences (both Pg = .03/.01, t_22_ < 1). Analyses of decision weights revealed that participants successfully focused on target signals and ignored concurrently presented distractors, as indicated by target weights which were around 9-times larger than concurrently presented distractors (.92 vs. .10, F_1,22_ = 1,693, p < .001, 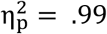). Critically, both target and distractor weights were statistically indistinguishable across low and high coherence conditions (main effect of coherence F1,22 < 1, coherence x stimulus type F_1,22_ = 1.34, p = .259). The control experiment showed that participants were able to discern target motion direction with comparable precision at both low and high coherence values, confirming that unbiased integration of the low and high coherence signals was, in principle, possible.

## Discussion

We have shown that behavioural decision weights and population-tuning profiles to average motion direction exhibit qualitatively similar biases in favour of stimuli of higher reliability. These findings demonstrate that temporal integration of discrete perceptual decisions is biased even when the bias is suboptimal, and when unbiased, optimal integration is possible. Our findings suggest that the brain encodes the reliability of sensory inputs, and that the encoded reliability is used automatically to weight respective inputs during integrative decision-making.

The relatively narrow distributions of error magnitudes and high response precision suggest that participants were able to successfully select target signals and ignore concurrently presented distractors. This finding was further corroborated by very low decision weights for distractors, which were five to nine times lower than the respective target weights. Perhaps most interestingly, the population-tuning modelling of distractor motion signals revealed no distractor-specific neural activity. As participants were given some preparation time (500–750 ms) before the motion onset, it is likely that this time was sufficient for attentional resources to be engaged exclusively on the target-motion stimulus. By contrast, motion decoding for target signals was robust and sustained well after signal offset, suggesting that the population-tuning model primarily captured decisionmaking processes as opposed to purely sensory-evoked activity patterns. With this in mind, it is likely that decoding of the average motion direction also reflected the dynamics of integrated decision making. One might ask whether it is possible that tuning to the average motion direction simply reflects tuning to the second target, given that the two directions were not entirely uncorrelated. This seems unlikely, however, because comparable tuning to the average motion direction should also have been observed in the first epoch (for which target signals were likewise not uncorrelated with the average), but this was clearly not the case. Moreover, if tuning to the average motion direction was driven by the second target then the time course of average motion tuning should have been similar to that of the second target, which, again, clearly was not the case. We therefore conclude that the robust tuning to the average target motion reflects the temporal dynamics of integrative decision making.

Unexpectedly, we observed a reliable order bias in behavioural responses, with larger decision weights for the first epoch than the second. Additionally, the colour-locked CPP had a steeper slope and a higher peak in the first epoch than the second. By contrast, the motion-locked CPPs did not differ much between epochs, at least prior to the peak of the CPP response. These findings suggest that the order bias originates from processes related to selecting the task-relevant dot patch, rather than from processing of the motion signals themselves. Perhaps most compellingly, the order bias was also evident in tuning to the average motion direction, with robust shifts in favour of the first target-motion direction. This order effect was independent of reliability-weighted source integration, as we observed the shift in population tuning for epochs that were matched for coherence. Additionally, as the stimuli across the two epochs were matched for low-level properties, the order bias cannot be stimulus-specific. Finally, the first-target (primacy) bias speaks against a simple memory decay explanation, which instead would predict a recency effect [29,30]. Taken together, the most parsimonious explanation for the primacy bias is that the effectiveness of attentional selection decreases from the first epoch to the second. Characterising attentional selection dynamics in relation to integrated decision-making was not the focus of the present study, and follow-up studies will be needed to address this issue in more detail. For the interested reader, we have recently conducted a study [37] that focused on the relationship between selective attention and decision-making using a similar experimental paradigm but a different analytical approach.

The reproduction task we employed enabled us to probe the nature of the representations underlying integrated decision-making. In typical decision-making paradigms [4], the response is a categorical decision, for example, whether motion direction is to the left or to the right. While forced-choice paradigms lend themselves to speeded responding, and permit the use of computational modelling to characterise different aspects of decision making, they do not capture the precision of the sensory and memory representations that underlie evidence accumulation processes. A well-known property of the brain’s responses to sensory input [18,38–40] is that they are graded, forming a probabilistic stimulus representation in feature space. In the case of motion signals, the large-scale neural representation of a given motion direction should resemble a bell-shaped curve with a peak over the actual direction which gradually decreases for motion directions further away from the peak. Classical decision-making paradigms would be sensitive to the location of the peak, but would have difficulty characterising the variability of the probabilistic representation – and that variability seems to play a critical role in integrated decision-making. Using mixture distribution modelling for behavioural measures, and population tuning modelling for neural measures, we were able to characterise both the peak and the variance of the underlying probabilistic representations.

A key question regarding reliability-weighted integrated decision-making concerns how two discrete representations get combined in support of an integrated decision. Previous research on signal integration on initial sensory encoding [23,24] and later decision-making [17,22,41] stages has suggested that a simple multiplication of two probabilistic representations could drive reliability-weighted integration. The multiplication, however, predicts *lower* variability [18,23] of the integrated representation relative to the variability of individual sources. Alternatively, the two discrete representations could be summed, rather than multiplied. To test the integration mechanism, we compared the variance of error magnitudes in the averaging task (main experiment) and the simple linear sum of error magnitude variances in the reproduction task (control experiment). For all combinations of dot coherence across the two epochs, the variance in the averaging task did not differ significantly from the sum of variance in the reproduction task (all p_FDR-corrected_ > .05). Therefore, it appears that for tasks such as the one employed here the two probabilistic representations are summed rather than multiplied.

The summation process appears to be biased, with larger weights for high-reliability representations. One possibility is that the weights directly reflect the *variability* of the probabilistic signal representations: a simple combination of two probabilistic representations of differing variances would be shifted in favour of the representation of higher reliability. Another possibility is that the weights reflect a *belief* about the accuracy of the respective representations. In this scenario, even though the two reliability levels afforded comparable accuracy (as confirmed in the control experiment), the strong perceptual differences between low- and high-reliability signals would have resulted in different beliefs about signals of different coherence. While at present we cannot adjudicate between the two potential mechanisms, the absence of strong coherence effects in our study suggests that the representations of the high-and low-reliability signals were comparably accurate, speaking in favour of the latter, *beliefs-as-weights* alternative. Further studies, most likely in combination with hierarchical computational modelling [42], would be necessary to address this issue conclusively. At present, computational models of decision-making in reproduction tasks are just beginning to appear [43,44], and more research will be needed before applying these models to integrative decision-making tasks.

In summary, here we have shown that combining two discrete, temporally separated signals in support of a single, integrated decision is biased in favour of higher reliability signals. Unlike previous studies in which reliability-weighted integration was statistically optimal, in the present study biased integration was suboptimal. These findings suggest that reliability-weighted integrated decision-making is automatic, taking place even when it is detrimental for performance.

## Materials and Methods

### Participants

26 neurotypical adult humans (mean age 21 years, 15 females) took part in the main experiment and another group of 24 neurotypical adults (mean age 22 years, 14 females) participated in the control experiment. All had normal or corrected-to-normal visual acuity and normal colour vision confirmed by Ishihara colour plates. The sample size was selected to achieve high power (*β* = .9 at *α* = .05) to detect a medium to large effect size (Cohen’s *d_z_* = .65) for a one-tailed, one-sample t-test between response error magnitude for low-and high-motion coherence. Based on behavioural performance and the EEG signals, four participants were identified as outliers in the main experiment (see below). Based on behavioural performance, one was identified as an outlier in the control experiment. The final sample comprised 22 and 23 participants in the main and control experiments, respectively. The study was approved by the Human Research Ethics Committee of The University of Queensland (approval nr 2016001247), and was conducted in accordance with the Human Subjects Guidelines of the Declaration of Helsinki. All participants provided written informed consent prior to experimental testing.

### Stimuli, task, and procedure

In both experiments, every trial started with a coloured cue, indicating the target colour (Fig. 1a). Two out of three easily discernible colours (pink, HSL values of 0, 75, 50; yellow, 90, 75, 50; and cyan, 270, 75, 50) served as target and distractor colours. The target-distractor colour pairs (e.g., pink target and blue distractor) were fixed per participant and counterbalanced between participants. After the cue, a circular patch (15.6 dva diameter) of 160 grey, randomly moving dots (.6 dva diameter, speed 2.5 dva/s, infinite dot life) appeared. To prevent a stimulus onset-evoked response from influencing electrophysiological measures of decisionmaking, the grey patch remained on screen for 1 s. Thereafter, the dot saturation increased gradually (over .25 s), revealing two overlapping patches of coloured dots (80 dots per patch) in the target and distractor colours. The colour saturation remained at maximum for 1 s and then gradually returned to grey. During maximum saturation, coherent motion signals were presented briefly (.5 s) in both patches. The onset of coherent motion was jittered (.25–.5 s) relative to the maximum colour saturation. The motion coherence was pseudo-randomly selected for every epoch of coloured dots, with low (40%) and high (80%) coherences presented equally often. Participants had to monitor for target motion signals and ignore distractors. A feedback stimulus was presented after every trial indicating response accuracy in that trial. Response accuracy rather than speed was emphasised, and participants were given ample time to respond (max. 6 s).

In the main experiment, two epochs of coloured dots were presented in every trial and participants had to reproduce the average motion direction of the two target signals while ignoring distractor motion events. The target motion in the first epoch was selected randomly from 0–360 degrees range in 1 degree steps. The target motion in the second epoch was selected from a ±30– 150 degree range relative to the first target. The distractor motion was within a ±30–150 degree range relative to the presented target motion. The dot coherence across the two epochs was selected pseudo-randomly so that all four combinations (low/high in the first epoch × low/high in the second) were presented equally often. Both behavioural and electroencephalography data were recorded.

In the control experiment, only one epoch of coloured dots was presented per trial and participants had to reproduce the target motion direction by adjusting the orientation of a response dial. Per participant, two pairs of motion directions (i.e., four in total) were selected as target motions from a range of directions (0–360 degrees in 15-degree steps). Within a pair, the directions were 30 degrees apart from each other and the two pairs were 180 degrees apart (e.g., 15 and 45, 195 and 225 degrees). Different combinations of directions were counterbalanced across participants. Only behavioural data were recorded. All other details were as in the main experiment.

### Apparatus

The experiments were conducted in a dark, acoustically and electromagnetically shielded room. The stimuli were presented on a 24” monitor with 1920×1080 resolution and a refresh rate of 144 Hz. The experimental software was custom-coded in Python using the PsychoPy toolbox [45,46]. EEG signals were recorded using 64 Ag-AgCl electrodes (BioSemi ActiveTwo) arranged in the 10-20 layout, and sampled at 1,024 Hz.

### Behavioural analyses

To identify outlier participants, the distributions of error magnitudes (i.e., the angular difference between the response and the correct answer) were compared with a uniform distribution (i.e., pure guessing) using the Kolmogorov-Smirnov test. Participants for whom the probability of the null hypothesis (i.e., a uniform distribution of error magnitudes) exceeded .001 were removed from further analyses. The remaining distributions per experimental condition and per participant were fitted to a theoretical model [47], and responses were separated into noisy target responses and random guesses. To quantify decision weights, a multiple-regression (OLS) model with a term for each of the presented motion directions, expressed as complex numbers, was fitted to the responses, separately per participant and experimental condition. The absolute value of the resulting regression coefficients reflects the influence of each of the presented coherent motion signals on the response, i.e., its decision weight.

### EEG analyses

EEG signals were analysed using the MNE-Python toolbox [48]. The data were offline re-referenced to the average electrode, low-pass filtered at 99 Hz and notch-filtered at 50 Hz to eliminate line noise. The recorded signal was pre-processed using the FASTER algorithm for automated artefact rejection [49]. The pre-processed signal was down-sampled to 256 Hz, segmented into 4 s periods between the onset of the first epoch and the response-display onset, baseline-corrected relative to -.1–0 s pre-trial and linearly de-trended. Outlier trials and participants were identified using the FASTER algorithm and removed from further analyses.

Next, whole-trial time-traces were further segmented into shorter (.5 s) periods time-locked to the onset of colour-saturation increase and the onset of coherent motion in the first and second epoch, and baseline-corrected relative to .1–0 s pre-onset interval. To characterise the temporal dynamics of evidence accumulation, we quantified an ERP known as the central-parietal positivity (CPP). Previous research has shown that the time-course of the CPP closely resembles the timecourse of evidence accumulation: specifically, its amplitude builds up gradually, the build-up slope is proportional to stimulus quality, and it is observed even in the absence of overt responses [11,12,32]. Visual inspection of the ERP topographies revealed a positive deflection in a cluster of central-medial electrodes (FCz, C1, Cz, C2, and CPz) consistent with the CPP ERP – to improve signal-to-noise-ratio, the average of these electrodes was used in further analyses. Next, the CPP voltage per time-sample, trial and participant was submitted to a stepwise linear mixed-effects model with epoch (first/second) and motion coherence (low/high) per epoch as fixed effects, and participant as a random effect. A likelihood ratio test between models of higher and lower complexity was used to assess the significance of each main effect and interaction terms. To control for multiple comparisons, the p-values for all time-samples and all model terms were jointly corrected using the false discovery rate algorithm [50].

To recover feature-specific information about motion signals from the EEG signals (Fig. 6), we used a population tuning curve model [34–36]. To that end, the first and the second epochs (1 s segments time-locked to the onset of coherent motion) from all trials were concatenated (ca 700 segments per participant), temporally smoothed by convolving the time series with a Gaussian window (SD = 16 ms), shuffled and split into 10 testing sets 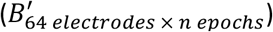. For each testing set, the remaining epochs were used as the training set (B). We modelled motion tuning as a set of 16 half-wave-rectified sinusoid channels raised to the 15^th^ power and centred on equidistant motion directions spanning 0–360°. The motion directions presented in each epoch were then convolved with the motion channels to yield a matrix of channel responses to motion signals (*C*_16 *channels* × *n epochs*_) in the training set. To model the mean EEG amplitude across training epochs, a fixed intercept term was added to the matrix of channel responses. The training EEG data were then modelled using the following linear model: *B* = *WC*, where W represents the weight matrix (*W*_64 *electrodes* x *16 channels*_) relating the EEG data and the population tuning model. The W matrix was estimated using ordinary least squares: *W* = *BC^T^*(*CC^T^*)^-1^ using Moore-Penrose inverse. Finally, the responses of the population tuning model in the testing set were estimated using the following equation: 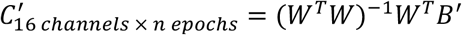. The profile of channel responses would reflect motion tuning: a uniform (i.e., flat) response profile would correspond to no tuning, whereas a prominent peak at channels close to the presented motion would reflect strong tuning.

**Figure 6.**
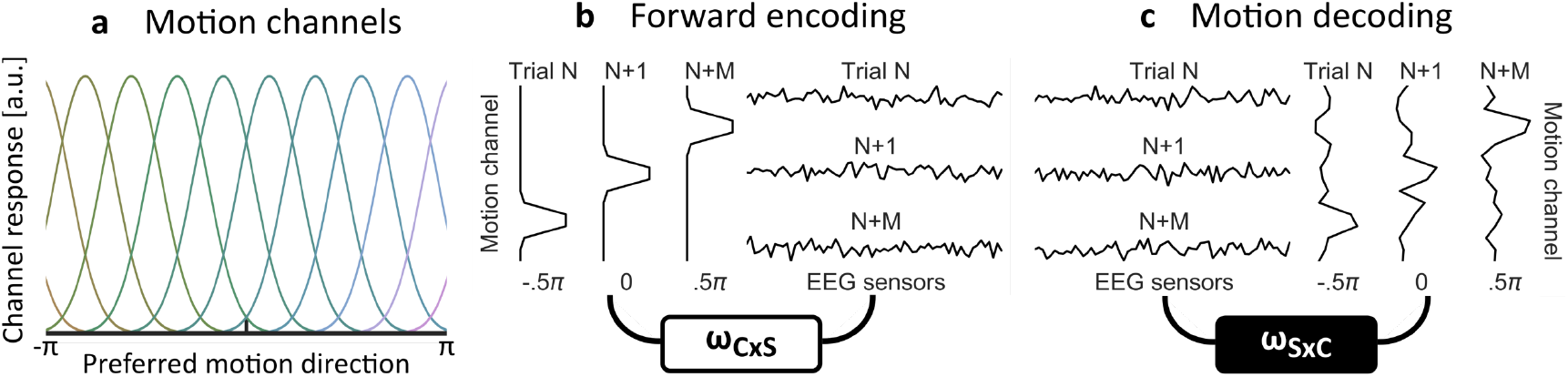
Illustration of the population-tuning analysis pipeline. (a) A set of hypothetical motion-selective channels is created, each of which is preferentially tuned to one of 16 equidistantly spaced motion directions spanning the full circle (-π to π). (b) The array of presented motion directions per trial in the training set of trials is convolved with the response profiles of the motion channels to yield trial-specific responses (left panel). The trial-specific responses are used as predictors to model the EEG time-traces from the respective trials (right). The forward encoding yields a weight matrix (ω_CxS_) relating channel responses to the EEG. (c) The inverted weight matrix (ω_SxC_) is applied to the EEG time-traces from the testing set of trials to retrieve the profiles of the motion channel responses.

The motion tuning analyses were conducted per time sample per participant. The tuning to target and distractor motion signals was analysed separately. We also estimated tuning to the average motion direction (i.e., the expected, average response) using the first and the second epoch data in two separate analyses. To characterise the overall tuning strength across different conditions, the vectors of channel responses were centred on the actual presented motion direction, temporally smoothed using a Gaussian window (SD = 16 ms) and averaged across trials. As an aggregate index of the response profile, we computed the dot product (*θ*) between the channel responses and the complex-valued preferred motion directions for respective channels. To quantify the tuning strength, we used the following equation: *f_θ_* = *abs*(*θ*)*cos*(*θ*). This equation yields a good, non-parametric descriptor of the overall tuning shape: (i) the *abs*(*θ*) yields 0 when there is no tuning (i.e., the distribution of channel responses is flat) and (ii) the *cos*(*θ*) reflects a mismatch between the expected preferred motion direction (which is normalized to 0) and the empirical peaks in the tuning profile. Similar to the ERP analyses, the tuning strength time series were analysed using stepwise mixed-effects general linear models, separately per time sample.

## Acknowledgements

This work was supported by the Australian Research Council (ARC, www.arc.gov.au) Centre of Excellence for Integrative Brain Function (ARC Centre Grant CE140100007). JBM was supported by an ARC Australian Laureate Fellowship (FL110100103). The funders had no role in study design, data collection and analysis, decision to publish, or preparation of the manuscript.

